# Anisotropic conductivity modeling for tDCS in Parkinson’s disease using multidimensional diffusion MRI

**DOI:** 10.64898/2026.08.02.742293

**Authors:** Santiago Osorio Jurado, Mikael Skorpil, Per Svenningsson, Rodrigo Moreno, Christoffer Olsson

**Affiliations:** Department of Biomedical Engineering, The City College of New York, City University of New York, New York, NY, USA; Department of Biomedical Engineering and Health Systems, KTH Royal Institute of Technology, Stockholm, Sweden; Department of Molecular Medicine and Surgery, Karolinska Institutet, Stockholm, Sweden; Department of Clinical Neuroscience, Karolinska Institutet, Stockholm, Sweden

**Keywords:** Transcranial direct current stimulation, Parkinson disease, Diffusion MRI, Electric field modeling, Finite element method, q-space trajectory imaging

## Abstract

Transcranial direct current stimulation (tDCS) dose depends on how brain conductivity is modeled. White matter anisotropy is conventionally estimated from single-shell diffusion tensor imaging (DTI). Multidimensional diffusion MRI (MD-dMRI), specifically q-space trajectory imaging (QTI), instead gives a mean tensor expected to carry less kurtosis bias. Our primary question was whether replacing the conventional single-shell tensor with this mean tensor would change the predicted field. We built, to our knowledge, the first MD-dMRI tDCS conductivity model and compared it against DTI and isotropic models in 29 participants (12 with Parkinson’s disease, 17 controls) across four montages, with the same mesh, electrodes, and solver. The three models agreed within a few percent. The two anisotropic models differed mainly in tensor orientation (about 21 degrees in white matter), with small differences in field magnitude. Field did not differ between patients and controls in any region or montage (which was an exploratory, underpowered comparison). Whole-brain electric field correlated with MR elastography stiffness (partial r = +0.58) but attenuated to non-significance once cerebrospinal fluid morphology was accounted for (r = +0.06 to +0.09). With no ground-truth field or conductivity available, the study establishes the feasibility of the MD-dMRI model and characterizes field sensitivity rather than improved dosimetry accuracy. The choice of diffusion tensor is second order for dose, which is primarily influenced by individual anatomy. For Parkinson’s disease, modeling efforts should focus on cerebrospinal fluid- and atrophy-aware head models and dose normalization, rather than a more complex diffusion tensor.

**Highlights:**

- Individual anatomy, more than the conductivity tensor, governs tDCS dose.
- A first tDCS head model from multidimensional diffusion MRI (QTI).
- MD-dMRI, single-shell DTI and isotropic fields agreed within a few percent.
- The anisotropic models differ mainly in orientation; their fields agree closely.

## 1. Introduction

Transcranial direct current stimulation (tDCS) is currently under investigation as a non-invasive therapy in Parkinson’s disease (PD), where it aims to modulate cortical and network excitability (Fregni et al., 2006; Broeder et al., 2015; Lefaucheur et al., 2017). Its effects depend on the electric field delivered to the target (Nitsche and Paulus, 2000), with a nonlinear dose-response (Jamil et al., 2017), yet the field cannot be measured directly and it varies substantially across individuals since it depends on head anatomy and on the electrical conductivity of brain tissue (Laakso et al., 2015; Huang et al., 2017). Because the dose-response relationship is nonlinear, an inaccurate field estimate can place a subject at the wrong point on that curve, which is one proposed contributor to the inconsistent clinical effects reported in tDCS trials for PD (Elsner et al., 2016). Personalized finite-element head models estimate the field from individual anatomy (Thielscher et al., 2015; Windhoff et al., 2013; Saturnino et al., 2019a).

White matter (WM) is electrically anisotropic, and its conductivity tensor is conventionally derived from single-shell diffusion tensor imaging (DTI) through a volume-normalized mapping, which translates the shape and orientation of the water diffusion tensor into the conductivity tensor (Tuch et al., 2001; Rullmann et al., 2009; Güllmar et al., 2010). However, the single-shell tensor is a biased estimate of the macroscopic tensor. A mono-exponential fit disregards non-Gaussian diffusion (Jensen et al., 2005), making the estimate dependent on the b-value and susceptible to errors caused by free water (Pasternak et al., 2009). Single-shell WM metrics vary widely across studies in PD (Atkinson-Clement et al., 2017), so the anisotropy derived from a DTI model is itself uncertain. Modeling a kurtosis term alongside diffusion yields a b-value-independent tensor estimate (Veraart et al., 2011).

Multidimensional diffusion MRI (MD-dMRI), in particular q-space trajectory imaging (QTI) with tensor-valued encoding, estimates the macroscopic mean tensor alongside the covariance of the intra-voxel tensor distribution, placing the non-Gaussian variance in the covariance term rather than in the mean (Westin et al., 2016; Topgaard, 2017; Szczepankiewicz et al., 2019). This extends the benefit of modeling the kurtosis separately, as in diffusional kurtosis imaging, to tensor-valued encoding. The resulting mean tensor is therefore expected to be less kurtosis-biased, and the decomposition has validated biophysical meaning and clinical relevance in tumor and WM disease (Szczepankiewicz et al., 2016; Nilsson et al., 2020; Andersen et al., 2020). Unlike conductivity tensor imaging, where conductivity is derived from an extracellular water diffusion tensor scaled by high-frequency electrical properties and volume fractions (Katoch et al., 2023), MD-dMRI has not, to our knowledge, been used to build a tDCS conductivity model, and it is unknown whether a less kurtosis-biased tensor changes the predicted field enough to matter for dosimetry.

Here, we build a tDCS conductivity model from the MD-dMRI mean tensor and ask whether replacing the conventional single-shell tensor with it changes the predicted field. Using the cohort of Olsson et al. (2025), 12 patients with PD and 17 controls scanned with both single-shell and MD-dMRI acquisitions, we compare three conductivity models, isotropic, single-shell DTI, and MD-dMRI, under an identical head model, electrode set, and solver. Our primary question is: for a given subject and montage, how much does the field change when only the conductivity model changes? As a secondary, exploratory question, we ask whether patients and controls differ in the modeled field. Additionally, we examine the field’s association with the brain’s mechanical properties, measured by magnetic resonance elastography (MRE) in the same cohort. In this study, we analyze the feasibility of building an MD-dMRI head model, and the sensitivity of tDCS dose to the choice of diffusion tensor.

## 2. Methods

### 2.1 Study design

We compared tDCS electric field predictions across three tissue conductivity models in one cohort of patients with PD and healthy controls (HC). The three models shared an identical head mesh, electrode configuration, and finite element solver, and differed only in how conductivity was assigned to brain tissue. The first model used a single isotropic value per tissue. The second, derived an anisotropic conductivity tensor from a conventional single-shell diffusion tensor acquisition. The third, derived the tensor from the macroscopic mean diffusion tensor estimated with MD-dMRI (QTI).

The study addressed two questions. The primary question was: for a given subject and montage, how much does the predicted field change when the conductivity model changes? Because every subject contributes to all three models, we treated this as a paired comparison. The secondary question was between groups: do patients and controls differ in the electric field metrics produced by using the MD-dMRI mean diffusion tensor? Since the cohort was not powered for this comparison, the results are exploratory. Figure 1 gives an overview of the modeling pipeline and the four electrode montages.

**Figure 1.**
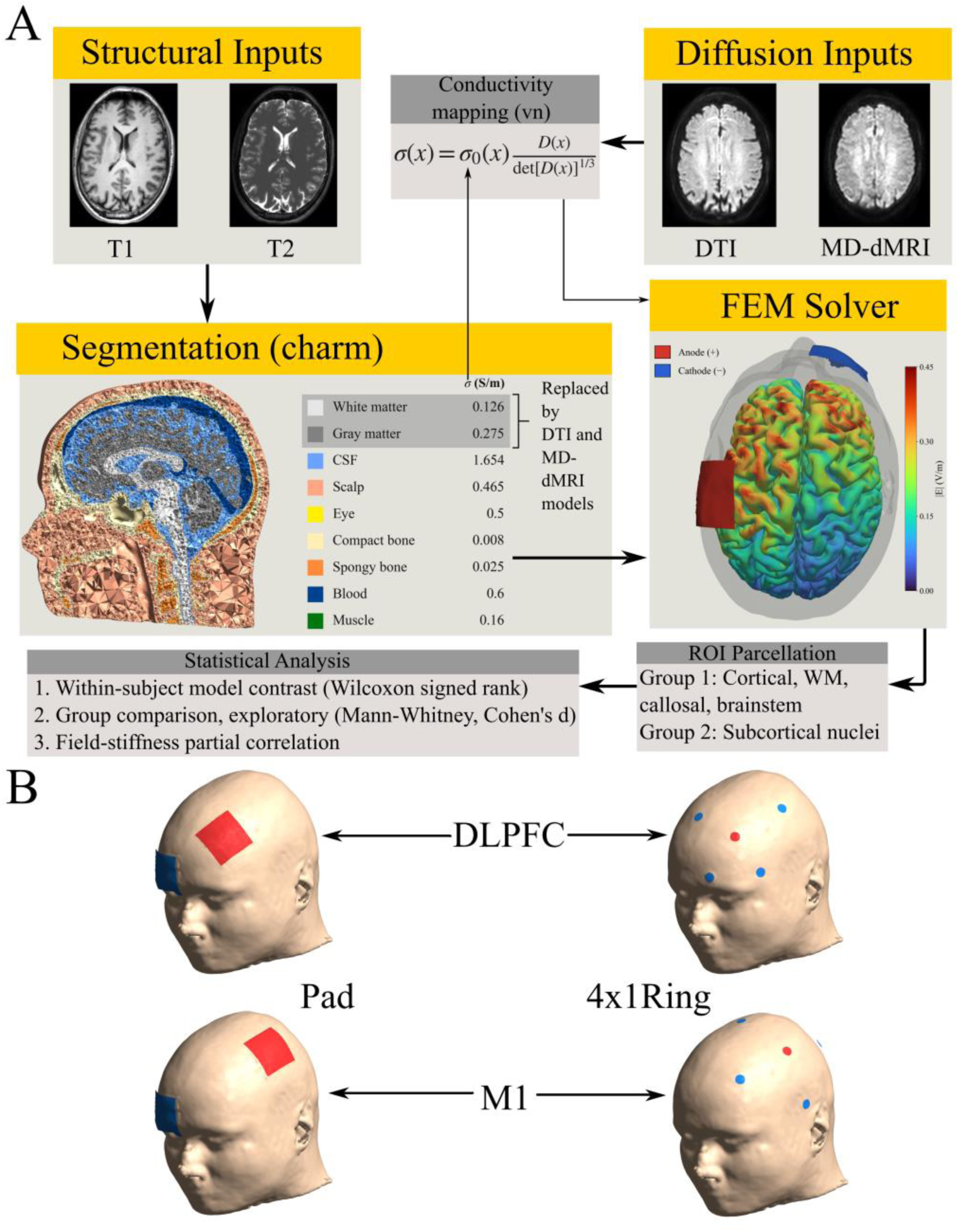
Overview of the modeling pipeline and electrode montages. (A) For each participant, the structural and diffusion MR images feed a single head segmentation and finite-element mesh. The three conductivity models (isotropic, DTI, and MD-dMRI) share that mesh and the same field solver and differ only in the conductivity assigned to brain tissue. (B) The four tDCS montages: conventional two-pad and 4 x 1 high-definition (HD) rings at the M1 (primary motor cortex) and DLPFC (dorsolateral prefrontal cortex) targets. In each head, the red electrode is the anode, and the blue electrodes are the cathodes.

### 2.2 Participants and data

We used the dataset described in Olsson et al. (2025). The cohort comprised 29 participants: 12 with PD, assessed in the ON medication state (3 women and 9 men; age 63 ± 9 years, range 47– 78), and 17 HC (3 women and 14 men; age 60 ± 6 years, range 50–78). The full demographic and clinical characterization is reported by Olsson et al. (2025). All participants gave written informed consent, and the original study was approved by the Swedish Ethical Review Authority (Dnr. 2022-03209-02). Our analysis was a secondary use of these data, covered by that approval. We analyzed the full cohort and excluded no participants.

All scans were acquired on a Philips Ingenia CX 3T system (Philips Healthcare, Best, the Netherlands) with a 32-channel head coil. The session provided a T1-weighted image (Turbo Field Echo, TR/TE 6.7/3 ms, 1 mm isotropic, full-head FOV), a T2-weighted image (TR/TE 3000/280 ms, 1 mm isotropic), a single-shell diffusion acquisition for the DTI model, and a multidimensional diffusion (QTI) acquisition for the MD-dMRI model. The QTI acquisition followed Topgaard (2017) and Szczepankiewicz et al. (2019), using linear and spherical b-tensor encoding at five b-values (0, 100, 700, 1400, 2000 s/mm²), TR/TE 4000/111 ms, 2.5 mm isotropic, in 6.5 minutes. The single-shell diffusion acquisition comprised 80 directions at b = 1500 s/mm² with one b = 0 volume, TR/TE 2658/83 ms, isotropic 2.0 mm resolution, multiband factor 3. This single-shell sequence was acquired in the same session but is not among the acquisitions reported by Olsson et al. (2025). The two diffusion acquisitions came from separate scans in the same session, so the DTI and MD-dMRI conductivity models come from separate acquisitions rather than two reconstructions of the same raw data.

### 2.3 Head model

We generated a tetrahedral head model for each subject with the charm pipeline in SimNIBS 4.6 (Thielscher et al., 2015; Puonti et al., 2020). Charm segments the T1- and T2-weighted images into tissue classes and produces a conforming finite element mesh. The electric field simulation assigns an isotropic conductivity to nine of these classes: WM, gray matter (GM), cerebrospinal fluid (CSF), scalp, eyes, compact bone, spongy bone, blood, and muscle. We ran charm with its default settings and inspected every segmentation visually. No manual corrections were applied.

### 2.4 Conductivity models

The three models differ only in the conductivity assigned to WM and GM. Every other compartment received the same isotropic conductivity in all three models, and the isotropic model used isotropic conductivities throughout. We used the SimNIBS 4.6 default scalar conductivities (shown in Figure 1), with WM and GM set to 0.126 and 0.275 S/m respectively (Wagner et al., 2004). The isotropic model serves as a baseline with no anisotropy. The primary comparison is between the two anisotropic models, with the conventional single-shell DTI tensor acting as the reference for the MD-dMRI tensor.

Both anisotropic models map a diffusion tensor to a conductivity tensor with the volume-normalized rule (Güllmar et al., 2010; Rullmann et al., 2009), implemented in SimNIBS as anisotropy_type = ‘vn’:

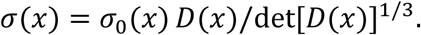

Here *D*(*x*) is the voxel diffusion tensor and *σ*_0_(*x*) the isotropic tissue conductivity. The conductivity tensor inherits the eigenvectors of *D*, and the determinant term fixes the geometric mean of its eigenvalues to one, so only the shape and orientation of *D* survive the mapping, and the scale of each tissue is set by *σ*_0_. We left both SimNIBS bounding parameters at their defaults from version 4.6, identical for both models.

For the DTI model, we fitted the single-shell diffusion tensor with FSL dtifit (Jenkinson et al., 2012) and passed it to dwi2cond (Opitz et al., 2011) for its T1 coregistration and reorientation, using a 12-degree-of-freedom (DOF) affine so the DTI model shares the registration class of the MD-dMRI model. Leaving dwi2cond otherwise at its defaults keeps the DTI model aligned with standard practice.

For the MD-dMRI model, the diffusion tensor was the QTI macroscopic mean tensor (Westin et al., 2016; Topgaard, 2017). We fitted the QTI signal with the diffusion tensor distribution covariance model, using the MD-dMRI framework (Nilsson et al., 2018), matching the processing applied to this cohort by Olsson et al. (2025). The QTI signal was spatially smoothed before fitting (isotropic Gaussian, *σ* = 0.7 voxels) to reduce noise and stabilize the covariance fit. The covariance model estimates more parameters per voxel than a single-shell tensor, and without smoothing it was non-positive-definite in about a fifth of brain voxels, with a noise-inflated WM anisotropy (near 15:1 in the most anisotropic voxels, above the 10:1 physiological ceiling, against near 4:1 when smoothed). The smoothing therefore trades spatial resolution for a physically plausible tensor. Its consequence for the between-model (MD-dMRI versus DTI) field magnitude comparison is taken up in Section 4.4. The DTI tensor received no comparable smoothing, so the two anisotropic inputs differ in resolution and smoothing as well as in estimation method. Therefore, the difference in electric field magnitude between the two models, measured as its 95^th^ percentile (*p*^95^ |E|; Section 2.8), cannot be attributed to the estimation method alone, as it is confounded by these acquisition and smoothing differences. Our claims about how the two tensor estimates differ rest instead on their orientation. The mean tensor is the first cumulant of that fit, the macroscopic mean diffusion tensor, estimated jointly with the covariance of the intra-voxel tensor distribution. Because the non-Gaussian variance is carried by the covariance term rather than absorbed into the mean, the mean tensor targets the same macroscopic tensor as the single-shell fit with reduced kurtosis bias (Westin et al., 2016; Veraart et al., 2011; Jensen et al., 2005). To obtain the MD-dMRI oriented macroscopic mean tensor required for conductivity mapping, we re-fitted the covariance model to the corrected linear and spherical encoding series, changing only the fit output used, not the upstream preprocessing. The encoding was carried through that motion and distortion correction, so the reconstructed orientation is in register with the corrected data. We then supplied that tensor to the same volume-normalized (‘vn’) mapping and solver used in the DTI model. The two anisotropic models are therefore identical in mapping and solver, differing only in the diffusion tensor and its acquisition. Voxels where the covariance fit returned a non-positive-definite or implausibly low-diffusivity mean tensor (a minority, concentrated at CSF borders and low-SNR edges) were set to isotropic before mapping, so they contribute neither spurious anisotropy nor orientation.

### 2.5 Registration and tensor reorientation

The conductivity tensors are defined in diffusion space and must be moved into the head model’s anatomical space before they can be assigned to mesh elements. The QTI diffusion data underwent distortion correction upstream using a synthetic reverse-phase-encode b = 0 (Synb0-DisCo; Schilling et al., 2019, 2020) followed by topup and eddy (Andersson et al., 2003; Andersson and Sotiropoulos, 2016). Since the data are already corrected for distortion, a 12-DOF affine is the appropriate registration class. In each subject, a nonlinear warp added spurious deformation without improving alignment. We registered the MD-dMRI model’s S0 image to the charm T2 with a 12-DOF affine (FSL FLIRT, mutual information; Jenkinson and Smith, 2001; Jenkinson et al., 2002), initialized from a rigid pre-alignment and rejected if degenerate, and applied that transform to every map.

Moving a tensor requires reorienting it, not merely resampling its components (Alexander et al., 2001). We warped the three eigenvalues independently as scalar maps (trilinear) and carried the orientation frame with vecreg, anchoring the principal axis to the registered principal eigenvector. After interpolation, the eigenvalue maps were sorted and paired once again with the frame by magnitude, so a boundary voxel where two maps cross cannot misassign an eigenvalue to an eigenvector. We validated each subject’s registration against the known fiber orientation of two reference structures segmented from each subject’s own anatomy: left-right across the corpus callosum and superior-inferior through the midbrain, which carries the cerebral peduncles. Each registration also passed automated field-of-view containment and S0-to-T1 edge-alignment checks and agreed with the fractional anisotropy (FA) registration to T1 provided with the cohort.

To check that the orientation difference between the two anisotropic models is not an artifact of aligning each diffusion scan to the head model separately, we computed the principal direction angle for each region using both tensors that were passed to the head model through the same final transform. For this control, the DTI tensor was first registered into the MD-dMRI native space, and both tensors were then taken to the head model by the one MD-dMRI S0-to-T2 affine. This shared-registration control is reported with the results.

### 2.6 Electrode montages

We simulated four montages, chosen to cover the two cortical targets most used in PD tDCS and to contrast conventional pads with focal high-definition arrays (Table 1). Two were conventional two-pad montages: a primary motor cortex montage (anode over C3, cathode over Fp2) and a dorsolateral prefrontal montage (anode over F3, cathode over Fp2), each with 5 x 5 cm pads at 2 mA. Two were 4 x 1 high-definition ring montages (Datta et al., 2009): a motor montage with the anode at C3 and cathodes at Cz, F3, T7, and P3, and a prefrontal montage with the anode at F3 and cathodes at Fp1, Fz, C3, and F7. In each ring montage, the anode delivered 2 mA and the four cathodes each carried -0.5 mA.

**Table 1.**
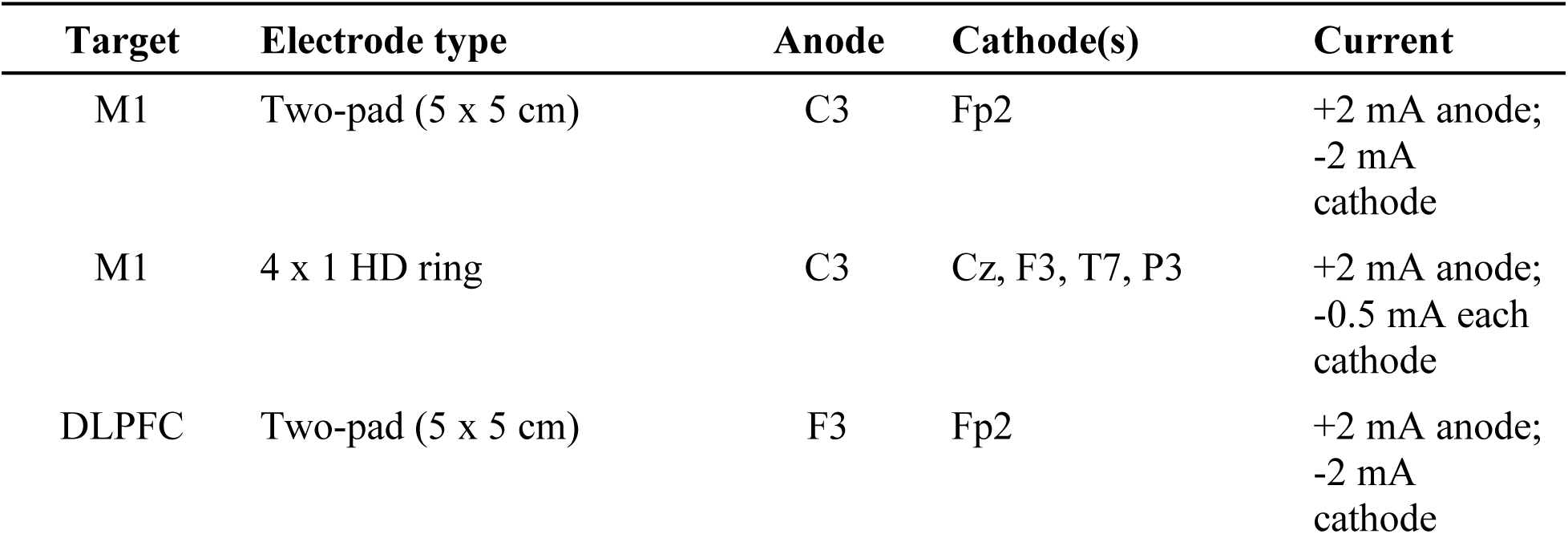
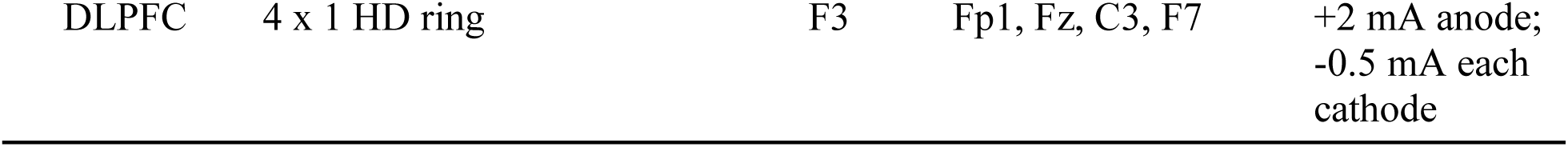
Electrode montages simulated. The four montages span two cortical targets (primary motor cortex, M1; dorsolateral prefrontal cortex, DLPFC) crossed with two electrode types: a conventional two-pad configuration (5 x 5 cm rectangular pads, 25 cm2, 0.080 mA/cm2 at 2 mA) and a 4 x 1 high-definition (HD) ring of 12 mm diameter disc electrodes. For each subject, all montages used the same head mesh, finite-element solver, and 2 mA total current, and the two electrode types at each target shared the anode position. Electrode labels are 10-10 EEG positions.

### 2.7 Regions of interest

We categorized regions into two groups based on their spatial resolution and role in the analysis. The primary group (Group 1) comprised large, well-resolved structures used for model comparison. The exploratory group (Group 2) comprised small subcortical nuclei at or near the 2.5 mm diffusion voxel, for which we report field exposure only.

Group 1 covered the cortical and WM lobes, the corpus callosum, the mesencephalon, and the pons. We parcellated cortex and WM with FreeSurfer 7.2 recon-all (Fischl, 2012; Desikan et al., 2006), grouping the Desikan-Killiany regions into frontal, parietal, temporal, and occipital lobes and taking the WM lobes from the wmparc segmentation. We segmented the brainstem with the Iglesias brainstem substructures module (Iglesias et al., 2015) and split out the mesencephalon and pons. These structures are large and well resolved, so they carry the primary field metrics.

Group 2 covered nine subcortical nuclei, reported bilaterally: the putamen, caudate, nucleus accumbens, external and internal globus pallidus, substantia nigra pars compacta and pars reticulata, red nucleus, and subthalamic nucleus, all taken from the CIT168 probabilistic atlas (Pauli et al., 2018) and warped to each subject with FSL FLIRT (atlas-to-MNI152-NLin6 affine) followed by SimNIBS mni2subject. The subthalamic nucleus is the principal target of deep brain stimulation in advanced PD (Deuschl et al., 2006), which makes its field exposure of interest. We treated this group as exploratory, reporting their electrical field metrics in the supplemental material (S-Table 2) over overlap-allowed masks, without comparison to any mechanical or microstructural measures. The larger nuclei are well resolved relative to the 2.5 mm diffusion voxel and overlap minimally, whereas the substantia nigra and subthalamic nucleus are at or below voxel size and subject to overlapping with other nuclei, so the latter are best read as field exposure estimates rather than precise values for each nucleus.

All regions of interest (ROIs) were defined anatomically, from each subject’s own FreeSurfer segmentation and atlas registration, and were fixed before the electric field was computed. No region was selected or adjusted based on the field it contained.

### 2.8 Electric field metrics and statistics

For each subject, montage, and conductivity model, we solved the electric field on the head mesh (Windhoff et al., 2013; Saturnino et al., 2019a) and summarized its magnitude within each ROI over the GM and WM elements. Specifically, we report two electric field metrics per region. The 95th percentile of |E| is the dose metric for the model comparison. It summarizes the regional field without being inflated by the outlier elements (Huang et al., 2017). The median of |E| is used for the age-adjusted group comparison and the stiffness correlation.

For the primary within-subject comparison, we tested in each region and montage whether the electric field metric differed between conductivity models with the Wilcoxon signed-rank test, using the matched-pairs rank-biserial correlation as the effect size (Kerby, 2014). For each region, we also summarized the within-subject difference in *p*_95_ |E| (MD-dMRI minus DTI) at the cohort median, with a 95% confidence interval from a percentile bootstrap (10000 subject-level resamples; Table 2). For the exploratory comparison between groups, we compared patients and controls under the MD-dMRI model. Following Olsson et al. (2025), we adjusted each field metric for a linear age effect by ordinary least squares and compared the residuals between groups with the Mann-Whitney U test, reporting Cohen’s d. We made no correction for sex, given the small number of women in each group. Our primary analysis consists of the 11 paired tests in the Group 1 regions under the M1 pad montage (*p*_95_ |E|). The other three montages and the median metric are secondary. The comparison between groups is exploratory. Additionally, we asked whether orientation divergence between the two anisotropic tensors drives the difference in the field between the models. Across all subjects and WM regions, where the principal direction is well defined, we correlated the V1 angle for each region with the absolute percentage difference between the two models, with reference to the DTI model, (Δ = 100 |*E*_MD-dMRI_ − *E*_DTI_| / *E*_DTI_) in *p*_95_ |E|. The V1 angle is the acute angle between the DTI and QTI mean tensor principal eigenvectors (*θ* = arccos(|*v*_DTI_ · *v*_QTI_|), the absolute value resolving the sign ambiguity), computed per voxel and reported as the cohort median of each subject’s ROI median. Before correlating, we centered each region on its mean across each subject, so the correlation captures the variation within ROI rather than differences between regions. We obtained the 95% confidence interval from a subject-clustered bootstrap that resampled the 29 subjects with replacement, since each subject contributes to several correlated regions.

**Table 2.** Region-wise paired electric field comparison, MD-dMRI versus DTI. , for the 11 Group 1 regions, M1 pad montage (anode C3, cathode Fp2, 2 mA), n = 29. Ctx, cortical lobe; WM, white-matter lobe. Δp_95_ |E| (V/m) is the within-subject difference in the 95th-percentile field magnitude over the gray and white matter elements of each region (MD-dMRI minus DTI), at the cohort median, with a percentile-bootstrap 95% confidence interval on that median. r_rb_ is the matched-pairs rank-biserial effect size (Wilcoxon signed-rank, paired); positive r_rb_ indicates MD-dMRI exceeds DTI, negative the opposite. q is the Benjamini-Hochberg FDR-adjusted p-value within the Group 1 family; q < 0.001 is shown as <0.001, and q < 0.05 indicates significance.

| Region | $\Delta p_{95} \vert E\vert $ (V/m) | 95% CI | $r_{rb}$ | $q$ |
| --- | --- | --- | --- | --- |
| Ctx Frontal | +0.0010 | [+0.0006, +0.0020] | +0.651 | 0.003 |
| Ctx Parietal | +0.0032 | [+0.0016, +0.0034] | +0.995 | <0.001 |
| Ctx Temporal | +0.0006 | [+0.0000, +0.0009] | +0.430 | 0.048 |
| Ctx Occipital | +0.0008 | [+0.0000, +0.0011] | +0.513 | 0.019 |
| WM Frontal | -0.0239 | [-0.0298, -0.0214] | -1.000 | <0.001 |
| WM Parietal | -0.0163 | [-0.0197, -0.0146] | -1.000 | <0.001 |
| WM Temporal | -0.0194 | [-0.0242, -0.0168] | -1.000 | <0.001 |
| WM Occipital | -0.0110 | [-0.0127, -0.0091] | -1.000 | <0.001 |
| Corpus callosum | -0.0053 | [-0.0077, -0.0008] | -0.586 | 0.007 |
| Mesencephalon | +0.0005 | [-0.0021, +0.0044] | -0.016 | 0.949 |
| Pons | -0.0050 | [-0.0078, -0.0018] | -0.862 | <0.001 |

The false discovery rate across regions was controlled through the Benjamini-Hochberg procedure within each montage, field metric, and test family (Benjamini and Hochberg, 1995), treating a Benjamini-Hochberg adjusted q < 0.05 as significant throughout. With n = 29 paired observations, a two-sided paired test at alpha = 0.05 has 80% power to detect a within-subject effect of *d_z_* about 0.54 (a noncentral-t approximation to the Wilcoxon signed-rank test). The between-group comparison, with 12 patients and 17 controls, has 80% power only for a group difference of about Cohen’s d = 1.1 and cannot exclude moderate effects, which is the second reason we treat it as exploratory.

Electric field computations were obtained using SimNIBS 4.6.0. Lobar and brainstem parcellations were taken from FreeSurfer 7.2 and subcortical nuclei from the CIT168 atlas. Diffusion preprocessing and registration used FSL 6.0.7.22. The QTI mean tensor was fitted with the MD-dMRI framework in MATLAB R2026a (The MathWorks, Natick, MA, USA). Statistical analyses were run in Python 3.11.14 with NumPy 2.3.0 (Harris et al., 2020) and SciPy 1.17.1 (Virtanen et al., 2020).

### 2.9 Association with mechanical properties

As a secondary analysis, we related the predicted field to the cohort’s mechanical measures, testing whether whole-brain median |E| covaried with MRE stiffness (|G*|, the magnitude of the complex shear modulus, computed whole-brain excluding ventricles and CSF) across all 29 subjects, following the correlation approach of Olsson et al. (2025). We computed the partial Pearson correlation between median |E| and stiffness, controlling for group (PD/HC) as a covariate, and adjusted the correlation p-values with the Benjamini-Hochberg false discovery rate procedure, reported as q throughout (Benjamini and Hochberg, 1995), as in the ROI analyses. Then, we added age as a second covariate, since both stiffness and head morphology change with age, and separately added CSF volume and the CSF fraction, computed as CSF volume divided by the sum of CSF, WM, and GM, both from the charm tissue segmentation. The CSF covariates test whether the field-stiffness relationship is driven by CSF morphology rather than a direct effect of tissue stiffness, since current shunts through the highly conductive CSF and less field reaches the cortex in subjects with larger CSF volumes (Indahlastari et al., 2020). Additionally, we computed the partial correlation of |E| with CSF fraction controlling for stiffness, and the collinearity between stiffness and CSF fraction, which we report descriptively, without false-discovery correction. The correlation was repeated under all three conductivity models and we report the MD-dMRI model as representative. Finally, as an input check, we correlated mean diffusivity with stiffness to confirm that our diffusion inputs reproduce the negative association reported by Olsson et al. (2025), whose stiffness maps we used directly. We did not report the stiffness-microstructure correlations as findings of this study, since they re-derive Olsson et al. on the same data.

## 3. Results

### 3.1 Participants

All 29 participants completed the structural, DTI, and MD-dMRI acquisitions and produced a head model that passed visual inspection of its segmentation. In every subject, the reoriented principal eigenvector agreed with the corpus callosum (left-right) and cerebral peduncle (superior-inferior) structural references.

### 3.2 Conductivity model comparison (MD-dMRI versus DTI)

The two anisotropic models produced very similar fields. Under the M1 montage the within-subject difference in the 95^th^ percentile field magnitude (*p*_95_ |*E*|) was small, only a few percent of the field, yet statistically resolvable in 10 of the 11 primary regions (Wilcoxon signed-rank, Benjamini-Hochberg-corrected within the primary group), and the difference changed sign with tissue type. In the cortical lobes, MD-dMRI slightly exceeded DTI, whereas in the WM lobes, corpus callosum, and pons, MD-dMRI was below DTI by a larger margin. The mesencephalon was the only region without a significant difference. The magnitudes and effect sizes per ROI are given in Table 2, and the between-model orientation difference is reported in Section 3.5.

The same sign pattern, MD-dMRI above DTI in cortex and below in WM, was consistent across all four montages, with one exception (HD-DLPFC occipital cortex; Figure 2; S-Table 1). Significance was reached in 9 of 11 for each of DLPFC, HD-M1, and HD-DLPFC montages.

**Figure 2.**
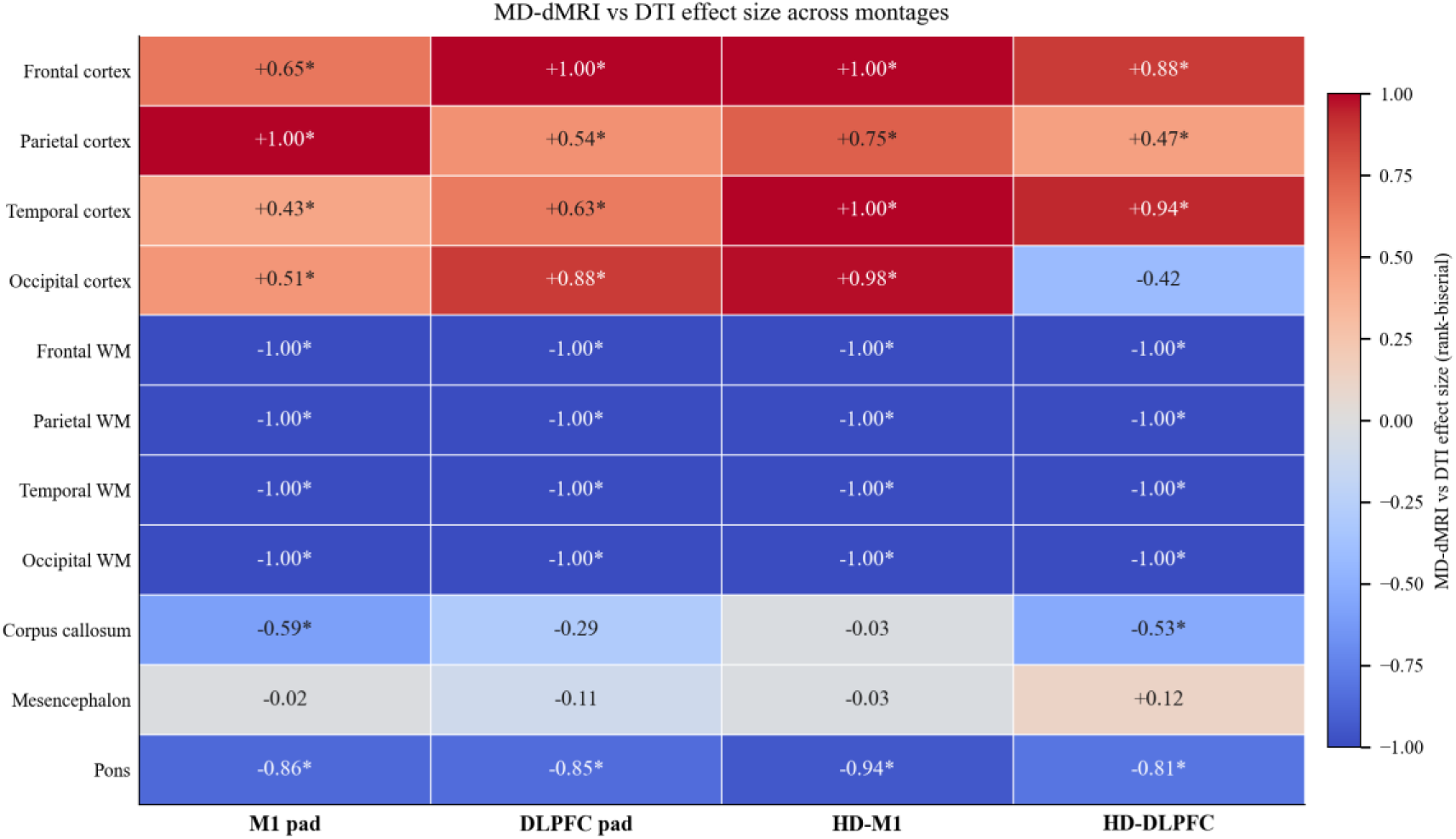
MD-dMRI versus DTI difference in the predicted field across montages. Each cell shows the matched-pairs rank-biserial (*r_rb_*) correlation for the within-subject difference in *p*_95_ |*E*| (MD-dMRI minus DTI), the effect size of the two-sided Wilcoxon signed-rank test. Rows are the 11 primary regions and columns the four montages (M1, DLPFC, HD-M1, HD-DLPFC), n = 29. P-values were adjusted with Benjamini-Hochberg control of the false discovery rate across the 11 regions within each montage; asterisks mark q < 0.05. The color scale is centered at zero (red, MD-dMRI above DTI; blue, MD-dMRI below DTI). Exact M1 values are given in Table 2.

### 3.3 Model difference relative to field magnitude

The absolute differences between models were small (Figure 3). In the cortical lobes, the three models lay within a few percent of one another. In the WM lobes and deep structures, the two anisotropic models sat slightly above the isotropic reference, with MD-dMRI below DTI, and the between-model differences stayed small relative to both the field magnitude and the between-subject interquartile spread. On the GM surface of a representative subject, the median absolute model difference was 0.0026 V/m, 1.8% of the median |*E*| (Figure 4).

**Figure 3.**
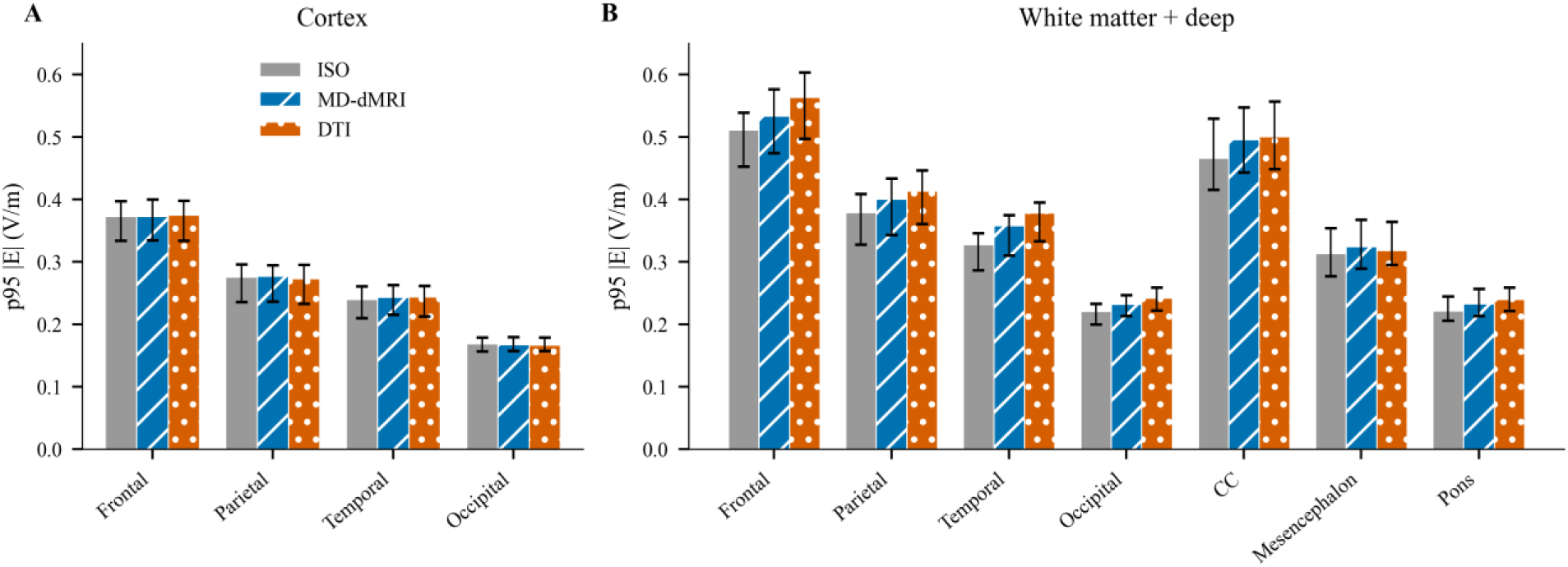
Predicted field magnitude for each conductivity model across regions, M1 montage. Cohort-median 95th-percentile |E| (V/m) over the GM and WM elements of each Group 1 region for the three conductivity models (ISO, MD-dMRI, DTI; n = 29); whiskers span the cohort interquartile range. (A) Cortical lobes. (B) WM lobes and deep structures (corpus callosum, mesencephalon, pons). The three models lie within a few percent of one another in the cortical lobes. In the WM lobes and deep structures, the anisotropic models sit slightly above the isotropic reference, with MD-dMRI below DTI. Regional effect sizes and significance, including the cortex versus WM sign difference, are in Figure 2 (all four montages) and Table 2 (M1).

**Figure 4.**
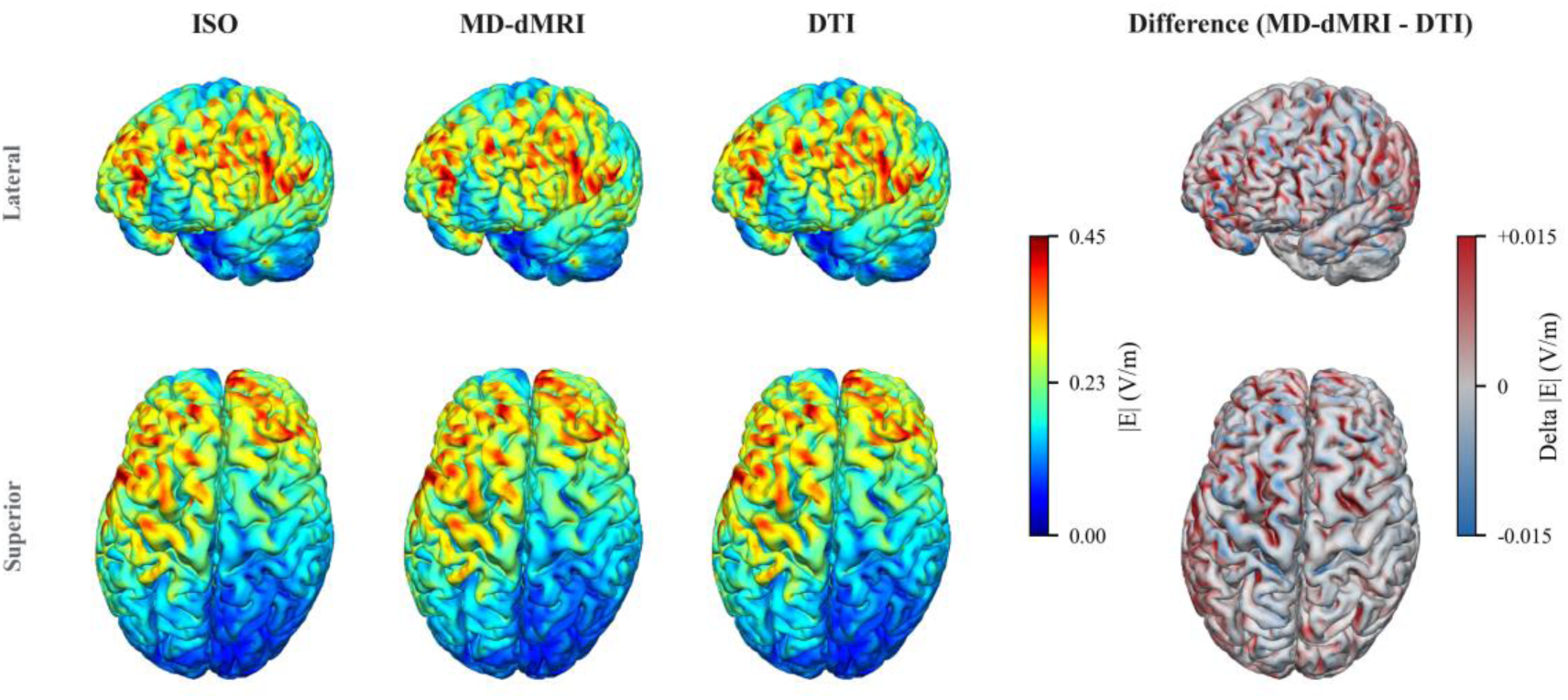
Predicted field on the cortical surface for each model, M1 montage, one representative participant. Columns show |*E*| for the isotropic (ISO), MD-dMRI, and DTI models and the difference between the two anisotropic models (MD-dMRI minus DTI), each at a lateral and a superior view of the GM surface. The ISO, MD-dMRI, and DTI columns share one color scale capped at 0.45 V/m (the 99.9th percentile of |*E*|). The difference column uses a separate symmetric scale whose cap (+/-0.015 V/m) is about 30 times smaller than the 0.45 V/m cap of the magnitude columns, so its colors should not be read as a large effect. The selected participant has a mean WM field difference between the two anisotropic models nearest to the cohort median.

### 3.4 Group comparison (PD vs HC)

Under the MD-dMRI model, patients and controls did not differ in field magnitude in any primary region under any montage (0 of 11 regions per montage; the largest effect in any region and montage was Cohen’s d = 0.63, minimum q = 0.48; age-adjusted Cohen’s d, Mann-Whitney on the age residuals, Benjamini-Hochberg-corrected; S-Figure 1). The null was consistent across all four montages (n = 29; 12 PD, 17 HC).

### 3.5 Orientation divergence between the two anisotropic models

The single-shell DTI and QTI mean tensors disagree mainly in orientation. Across the cohort, their principal directions differed by a median of about 21 degrees in the WM lobes and 13 to 18 degrees in the corpus callosum and brainstem, well above the near-zero angle expected if the two agreed. In GM, the angle rose to about 45 degrees, approaching the 60-degree random-axis chance level without reaching it, where the near-isotropic tensors leave the principal direction ill-defined (Figure 5). A single shared registration barely changed the divergence (22 degrees, against 21 under each tensor’s own registration; S-Table 3), so it comes from the tensor estimate, not from registering the two scans separately. The orientation difference did not, though, translate into a field difference: within regions, the V1 angle and the between-model change in *p*_95_ |E| were essentially uncorrelated (partial correlation controlling for region, r = -0.05, 95% subject-clustered bootstrap CI [-0.20, +0.12]; 29 subjects, 7 regions). We can make this claim for orientation but not for tensor shape: the shape and anisotropy difference between the models is confounded by the smoothing and native resolution mismatch of the two acquisitions (S-Table 3), so it is not a clean comparator.

**Figure 5.**
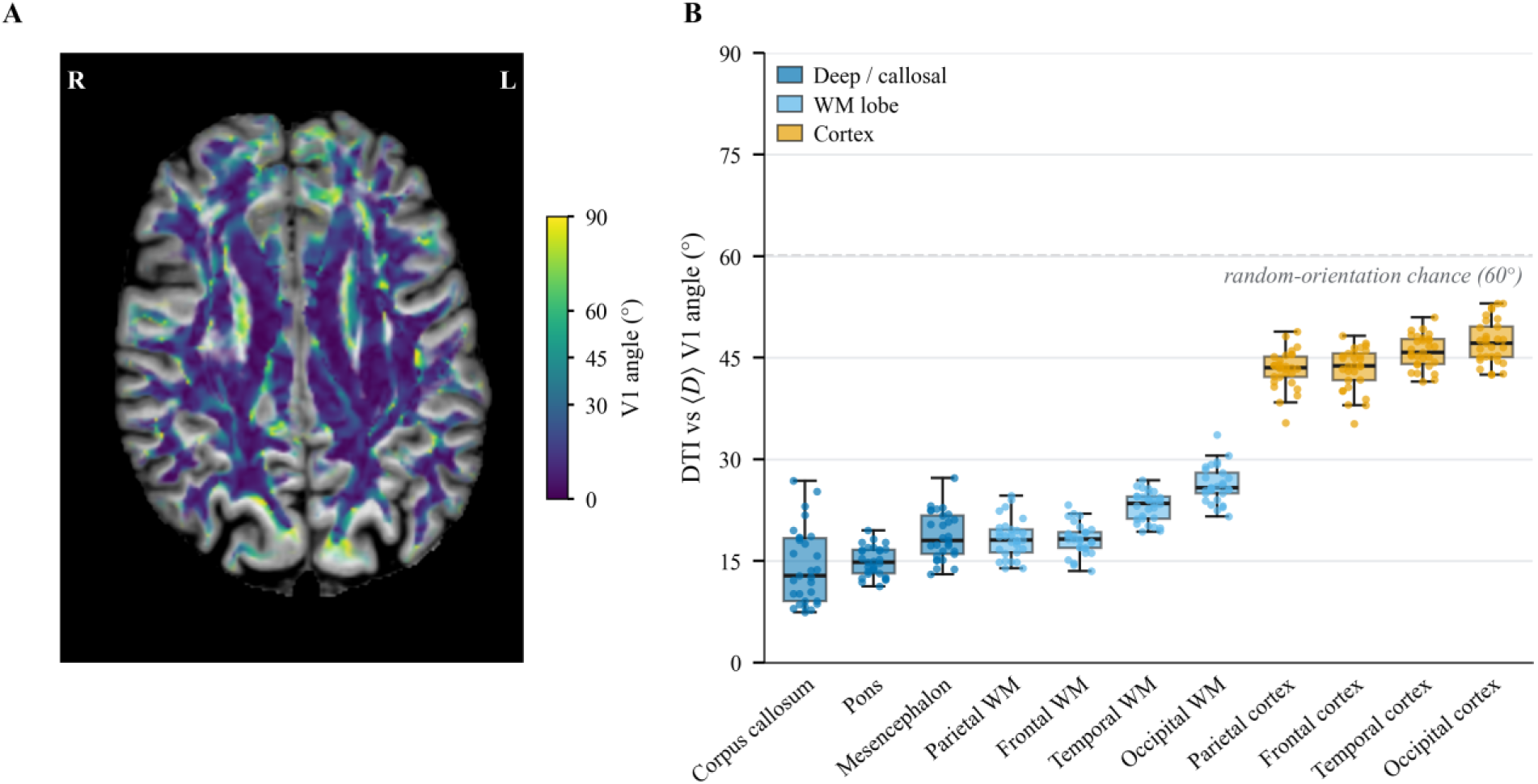
Orientation divergence between the two anisotropic models. Acute angle between the principal diffusion direction (V1) of the single-shell DTI tensor and that of the QTI mean tensor. A small angle means the two estimates point nearly the same way. (A) Voxel-wise angle on a representative transverse slice over a T1 underlay, across the WM and GM mask. Voxel opacity tracks fractional anisotropy, rising from zero at FA = 0.10 to full opacity at FA = 0.30, so strongly anisotropic voxels read solid and nearly isotropic ones fade into the underlay. (B) Angles across the cohort (n = 29): one box per region, ordered by median and colored by anisotropy regime (deep and callosal structures, WM lobes, cortical lobes), with individual subjects shown as points. The dashed line is the chance level for sign-ambiguous directions, the 60-degree median angle between two random three-dimensional axes. Values per region and the shared-registration control are in Results 3.5 and S-Table 3.

### 3.6 Association between predicted field and MRE stiffness

Whole-brain median |*E*| covaried positively with individual MRE stiffness, controlling for group (partial Pearson r = +0.58, q = 0.002; Figure 6A). The correlation was nearly identical across the three conductivity models (ISO, DTI, and MD-dMRI: r = +0.579 to +0.592, all q = 0.002), so the MD-dMRI model is shown as representative. Further adjusting for age left the association significant (partial r = +0.48, q = 0.018). Controlling instead for CSF volume attenuated it to non-significance (partial r = +0.09, q = 0.66). Expressing CSF as a head-size-normalized fraction rather than a volume, a sensitivity check, likewise left only a negligible partial correlation (r = +0.06). Examining the other way around, whole-brain |E| decreased with CSF fraction (r = -0.66; Figure 6B). Controlling for stiffness reduced this field-CSF correlation only partially (partial r = -0.41). Stiffness and CSF fraction were strongly collinear (r = -0.84). As an input check, mean diffusivity showed the expected negative correlation with MRE stiffness (Olsson et al., 2025). We do not report it as a finding, since it re-derives their results on the same data.

**Figure 6.**
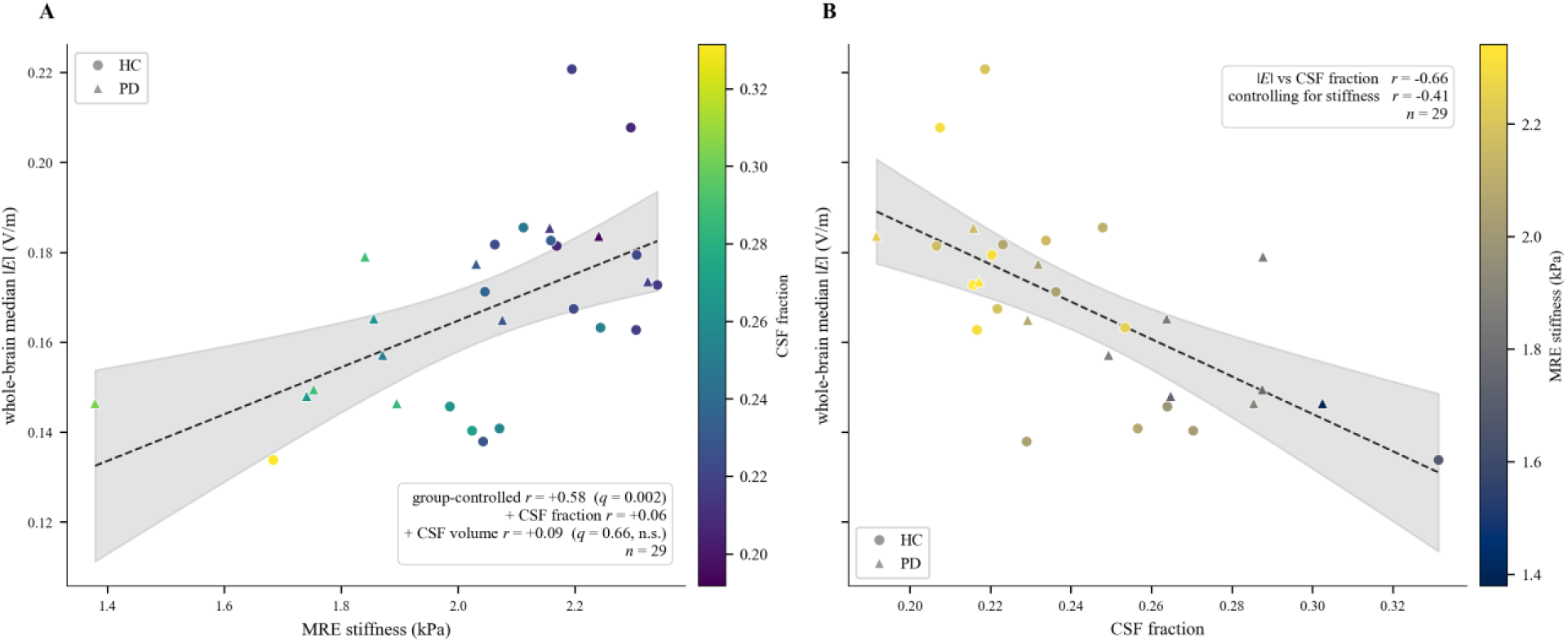
Association between the predicted field and MR elastography stiffness. (A) Whole-brain median |*E*| (MD-dMRI model, M1 montage) plotted against individual brain stiffness (kPa) for the 29 participants. Markers are colored by CSF fraction and shaped by group (circles, HC; triangles, PD). The dashed line is an ordinary-least-squares fit with its 95% confidence band. The partial Pearson correlation controlling for group is r = +0.58 (q = 0.002) and is near-identical across the ISO, DTI, and MD-dMRI models (r = +0.579 to +0.592). Controlling for CSF volume reduces the correlation to r = +0.09 (q = 0.66) and expressing CSF as a head-size-normalized fraction likewise reduces it to r = +0.06. (B) Whole-brain median |*E*| plotted against CSF fraction (r = -0.66), colored by stiffness, so each panel’s x-axis is the other panel’s color. Controlling for stiffness only reduces the association to r = -0.41.

## 4. Discussion

We built, to our knowledge, the first human tDCS conductivity model from the macroscopic mean tensor of MD-dMRI and compared it in a PD cohort against a conventional single-shell DTI tensor and an isotropic model, changing only the conductivity assigned to brain tissue while the head mesh, electrodes, and field solver stayed constant. The resulting field changed little: it depended on tissue type and was consistent across all four montages, but the change was small and stayed close to both the DTI and the isotropic model. Patients and controls did not differ significantly. The field correlated positively with brain stiffness, yet that association attenuated to non-significance once we controlled for CSF fraction, while the electrical field’s association with CSF fraction was more robust.

### 4.1 The choice of diffusion tensor is second order for dose

Because the multidimensional and single-shell tensors were passed through the same volume-normalized mapping and the same solver, the comparison isolates the effect of the tensor estimate on the field. Earlier work instead varied the conductivity mapping formula applied to a single diffusion tensor (Shahid et al., 2013), whereas here the mapping is held constant and only the tensor estimate changes. Such isolation preserves orientation, but not magnitude, because the two acquisitions differed in native resolution and smoothing, so the magnitude difference reflects that acquisition mismatch as much as the tensor estimate. This is also distinct from conductivity tensor imaging, which builds conductivity by scaling a conventional diffusion tensor with high-frequency electrical properties rather than from tensor-valued diffusion encoding (Katoch et al., 2023; Sajib et al., 2018; Mandija et al., 2026).

The difference we found was dependent on tissue type and consistent across montages, which lets us attribute it to the model used rather than the electrode montage. GM, which is close to isotropic, showed the three models converging. WM, where the tensor orientations genuinely diverge, showed a consistent field difference, on the order of a few percent, though the magnitude of that difference is confounded by acquisition and smoothing (see Section 4.4). This matches what recent work with the same segmentation and mapping tools reported: DTI-based anisotropy adds little to cortical transcranial electrical stimulation predictions, and it is the scalar tissue conductivities, not the anisotropy, that dominate tDCS modeling uncertainty (Schmidt et al., 2015; Saturnino et al., 2019b; Mosayebi-Samani et al., 2025). An earlier subject-specific analysis found that WM anisotropy changes the cortical surface field by only a few percent (Suh et al., 2012), though it reported a larger effect in deep brain regions, and realistic WM anisotropy likewise had only a marginal effect on EEG source localization (Lee et al., 2009). Estimates of the effect are not unanimous, and neglecting anisotropy altogether can shift the field by more than ten percent and its orientation by roughly twenty degrees (Caiani et al., 2026). Our comparison is a much narrower one since it is between two anisotropic tensor estimates rather than between anisotropic and isotropic models, and there the difference is small. Because the volume-normalized mapping keeps only the shape and orientation of the tensor, and because the electric field is only modestly sensitive to conductivity anisotropy (Opitz et al., 2011; Suh et al., 2012; Mosayebi-Samani et al., 2025), what separates the two models is orientation (Figure 5). The multidimensional mean tensor is expected to be the less kurtosis-biased, but without a measured conductivity, we cannot show which model is more accurate. What we can say is that changing the tensor changed the predicted dose only slightly, because the field is insensitive to this orientation difference at the present scale.

### 4.2 Individual anatomy, not the conductivity tensor, governs the dose

The link between the field and brain stiffness is consistent with a morphology effect. The whole-brain correlation was positive and did not depend on group, but it disappeared once we accounted for CSF, whether as total volume or as a fraction of brain plus CSF volume, and it was the same under all three conductivity models. CSF diverts injected current and lowers the field that reaches cortex (Huang et al., 2017), and its volume is one of the larger sources of between-subject variability in tDCS dose (Minjoli et al., 2017; Indahlastari et al., 2020). Individual head anatomy, CSF and skull thickness in particular, dominates that variability, to the point where stimulation with a fixed dose varies by more than a factor of two across people (Laakso et al., 2015; Evans et al., 2020), and the spread grows with more focal montages such as the high-definition arrays used here (Mikkonen et al., 2020). Brain stiffness falls with age, faster than volume does (Sack et al., 2011), and falls further in PD (Lipp et al., 2018; Olsson et al., 2025). Atrophy lowers stiffness and enlarges CSF compartments, which is why stiffness and CSF fraction were strongly collinear in this cohort (Section 3.6). Taking CSF into account eliminated the field-stiffness correlation, whereas adjusting for stiffness only partly reduced the field-CSF fraction correlation. Since CSF diverts injected current and stiffness has no direct electrical effect on the field, this is the expected result. Stiffness appears to track the same morphology that shapes the field rather than acting on the field through conductivity, which would explain why the association was the same under all three conductivity models.

We found no evidence of a group difference in the field in any region or montage. The comparison was underpowered at 29 participants, and we treat it as exploratory, but a null is also what the small model effect predicts: if disease changes the diffusion tensor only slightly as far as the field is concerned, the field will not separate patients from controls. Where predicted fields do separate patients from controls, it is generally due to gross macrostructural changes, whether the focal lesions of stroke (Kimura et al., 2026) or the anatomical differences of other clinical cohorts, in which a lower field was observed in patients than controls (Mizutani-Tiebel et al., 2022). PD in this cohort is not associated with macrostructural changes of that magnitude, so a group difference would not be expected on anatomical grounds. Thus, we interpret this null as consistent with subtle anatomy. For tDCS in PD, where the therapy shifts cortical excitability in a dose-dependent way with the delivered field (Nitsche and Paulus, 2000; Jamil et al., 2017) yet stays clinically modest and variable (Benninger et al., 2010; Broeder et al., 2015; Lefaucheur et al., 2017; Elsner et al., 2016), the individualization that matters is anatomical. A standard single-shell diffusion tensor already captures the anisotropy that reaches the dose, so modeling efforts are better spent on accurate CSF and atrophy segmentation, or on field-based dose normalization (Evans et al., 2020), rather than on refining the diffusion tensor.

### 4.3 Field exposure of the deep targets

A more accurate orientation should matter most in deep WM near stimulation targets. Transcranial current reaches subcortical structures (Datta et al., 2013), and our deep readouts show measurable field in the subcortical nuclei under all four montages (S-Table 2). The subthalamic nucleus and internal globus pallidus are the principal effective deep brain stimulation targets in advanced disease (Deuschl et al., 2006; Mao et al., 2019), so their exposure is worth knowing. These readouts are exploratory and resolution-limited, with the midbrain and subthalamic estimates indicative rather than precise, so we raise the deep target case as a reason to acquire higher resolution, not as a finding.

### 4.4 Limitations

This study has several limitations. The two anisotropic models came from separate diffusion scans that differed in resolution and smoothing, so the WM part of the magnitude difference is partly a preprocessing artifact rather than a property of the tensors. The smoothing that stabilizes the covariance fit also lowers its WM anisotropy, and removing it does not fix the comparison, because the unsmoothed fit is inflated by noise and non-positive-definite in about a fifth of voxels. Neither setting yields an unbiased magnitude, so we do not read the magnitude difference as an effect of the tensor choice. Instead, we lean on orientation. Because the two scans were separate, each tensor was aligned to the head model by its own 12-DOF affine. This does not bias the within-subject field comparison, where each tensor is placed correctly in the subject’s own anatomy. The shared-registration control (Section 3.5) shows that this orientation difference is not attributable to registration alone. It may, however, partly reflect angular sampling. The DTI acquisition used 80 diffusion directions, whereas the linear encoding of the MD-dMRI acquisition sampled 6 to 21 directions per shell. The mean tensor is therefore expected to be less kurtosis-biased but more affected by the lower angular resolution of its acquisition, and we interpret the orientation divergence as a difference between the tensor estimates rather than as evidence that either orientation is more accurate. The cohort is small, and predominantly male (see Olsson et al., 2025), which limits both the power of the group comparison and sex-stratified generalizability. The stiffness association is cross-sectional, and its covariates (CSF volume, age, and stiffness) are collinear, so even though the field’s association is with CSF rather than stiffness, this is still consistent with a shared morphology driver, not proof of a mediation pathway. The study is computational, with no measured field to validate against. Intracranial recordings are themselves too noisy to adjudicate modeling choices such as anisotropy (Puonti et al., 2019), but non-invasive magnetic resonance current density imaging provides a forward route (Göksu et al., 2018). Against these, the within-subject comparison is internally stable. The WM lobe difference is unanimous in sign and so does not hinge on the multiple comparison correction, but because its magnitude is confounded by preprocessing, we do not read it as a tensor effect. The tensor-estimate claim rests on the orientation divergence, and the agreement between the three models on field magnitude and the collapse of the stiffness association under CSF control are further checks that do not lean on any single modeling choice.

### 4.5 Future directions

Several steps would sharpen the comparison. Matching the two diffusion acquisitions in resolution and smoothing would separate the tensor effect from preprocessing and let the magnitude difference be read directly. A larger, prospectively recruited cohort would give the group comparison real power, and measured fields would test whether a better orientation provides a more accurate dose. A single macroscopic tensor cannot separate microscopic anisotropy from orientation dispersion, so it understates conductivity anisotropy where fibers cross or disperse. MD-dMRI separates the two and, through the covariance of the intra-voxel tensor distribution, resolves microstructure that a single mean tensor sets aside (Westin et al., 2016; Topgaard, 2017; Lampinen et al., 2017). Building a conductivity model on this richer microstructure is a natural next step, though because these microscopic measures have no orientation of their own (Westin et al., 2016), such a model would have to combine them with a fiber orientation distribution rather than read a conductivity tensor from them directly, and it would need an acquisition able to estimate the intra-voxel diffusion variance reliably (Szczepankiewicz et al., 2019). Combined with direct validation against a measured conductivity, such a model could reflect the electrical properties of WM more accurately than a single macroscopic tensor allows.

## 5. Conclusion

We showed that a tDCS conductivity model can be built from MD-dMRI, and that its predictions stay close to those of the DTI and isotropic models in PD. The model choice shifted the predicted field in a tissue-dependent way, but the shift was small, and what we can attribute to the tensor estimate is a difference in orientation, not in magnitude, since the magnitude difference is confounded by acquisition. Patients and controls showed no detectable difference in this underpowered, exploratory comparison. For tDCS dosimetry in PD, then, it is individual anatomy, more than the choice of diffusion tensor, that governs the dose, so individualized dosing is better served by CSF- and atrophy-aware head modeling than by a more elaborate diffusion tensor. The value of MD-dMRI may ultimately lie less in the mean tensor than in the microstructure it also resolves. Whether that can yield a more accurate conductivity model is an open question, and one that would need to be validated with measured conductivity.

## Data and code availability

The MR imaging data analyzed here (the cohort of Olsson et al., 2025) are subject to ethical and legal restrictions and cannot be shared publicly. Anonymized data can be made available to researchers who meet the applicable legal and ethical requirements, on reasonable request to the corresponding author. The analysis and modeling code is openly available at https://github.com/santioso-hue/MRE-tDCS-PD.

## Author contributions

**Santiago Osorio Jurado:** Conceptualization, Methodology, Software, Formal analysis, Investigation, Visualization, Writing – original draft, Writing – review & editing. **Mikael Skorpil:** Resources, Writing – review & editing. **Per Svenningsson:** Resources, Writing – review & editing. **Rodrigo Moreno:** Supervision, Resources, Writing – review & editing. **Christoffer Olsson:** Supervision, Data curation, Resources, Writing – review & editing.

## Acknowledgments

The acquisition and preprocessing of the diffusion and magnetic resonance elastography data reused in this study were supported by Hälsa, Medicin och Teknik (grant FoUI-992049), MedTechLabs, Digital Futures, Vinnova through AIDA (project 2319), and the Swedish Research Council (grant 2022-03389). S.O.J. was partially supported by the International Research Experiences for Students (IRES) program through the Chemical Engineering Department Travel Fund at The City College of New York. The funding sources had no role in the study design, in the analysis or interpretation of the data, or in the preparation of this article. We would like to thank the clinical research scientists at Philips and the MR physicists at Karolinska University Hospital in Huddinge, who set up the imaging protocols and supported the postprocessing of the original cohort study.

## Supplementary material

- **S-Figure 1.** PD versus HC comparison of the predicted field across montages (exploratory): age-adjusted Cohen’s d for MD-dMRI median |*E*| in the 11 primary regions under each montage.
- **S-Table 1.** All montage region-wise paired electric field comparison, MD-dMRI versus DTI.
- **S-Table 2.** All montage predicted electric field exposure of the Group-2 deep nuclei (MD-dMRI model).
- **S-Table 3.** Per-region principal-direction (V1) angle between the DTI and QTI mean tensors, under each tensor’s own registration and under a single shared registration.

