## Supplement for "Anisotropic conductivity modeling for tDCS in Parkinson’s disease using multidimensional diffusion MRI"

### Supplementary material

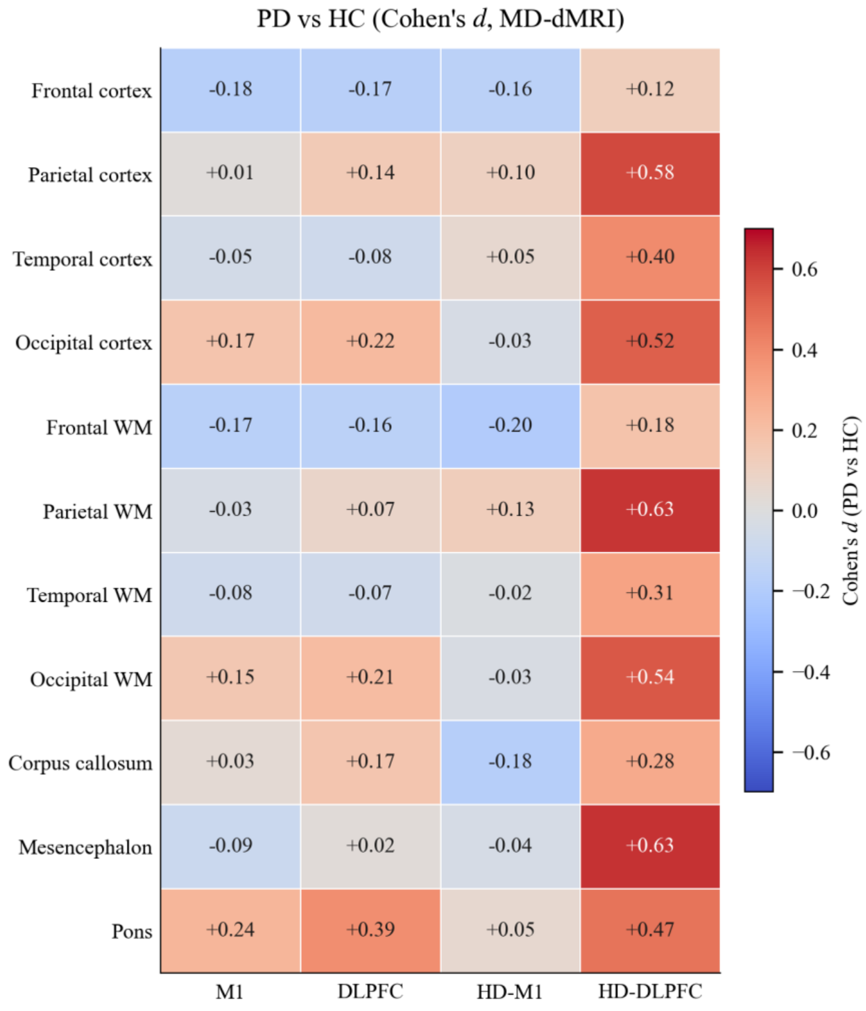


BH $q < 0.05$: 0/44 across all montages

**S-Figure 1.** Exploratory PD-versus-HC electric-field group differences across montages. Per-ROI age-adjusted Cohen's d (PD vs HC) for the MD-dMRI median |E| under each of the four montages (M1, DLPFC, HD-M1, HD-DLPFC; columns) across the 11 Group-1 ROIs (rows). Each ROI's per-subject median |E| is residualized on age by ordinary least squares before Cohen's d (PD residuals vs HC residuals); two-sample tests on the residuals are Benjamini-Hochberg corrected across the 11 ROIs per montage (* = q < 0.05). No region reached significance in any montage (0/11 per montage), and the null is consistent across all four, indicating that the MRE brain-stiffness group differences reported for this cohort are not reflected in a detectable electric-field group difference. The comparison is exploratory and underpowered (n = 29; 12 PD, 17 HC).

**S-Table 1.** All-montage region-wise paired electric-field comparison, MD-dMRI versus DTI, for the 11 Group 1 regions (four cortical lobes, four white-matter lobes, corpus callosum, mesencephalon, pons) under each of the four montages (M1 and DLPFC pad montages; HD-M1 and HD-DLPFC high-definition montages), n = 29. The M1 block reproduces Table 2. Ctx, cortical lobe; WM, white-matter lobe; CC, corpus callosum. ISO, DTI, and MD-dMRI are the cohort-median within-subject 95th-percentile electric-field magnitude ($p_{95}$ $|E|$, V/m) over the gray-matter and white-matter elements of each region, for the isotropic, single-shell DTI, and MD-dMRI conductivity models. Delta (V/m) is the within-subject paired difference in $p_{95}$ $|E|$ (MD-dMRI minus DTI), computed per subject and reported at the cohort median, with a percentile bootstrap 95 percent confidence interval (10000 resamples, fixed seed) on that median. $r_{rb}$ is the matched-pairs rank-biserial effect size for MD-dMRI versus DTI (two-sided Wilcoxon signed-rank, paired across subjects); positive $r_{rb}$ indicates MD-dMRI exceeds DTI across paired subjects and negative $r_{rb}$ indicates the opposite, and the sign of the paired Delta and $r_{rb}$ reflects the within-subject difference direction. Because the median is not a linear operator, the ISO, DTI, and MD-dMRI columns are marginal cohort medians and are not the terms of the paired difference: in near-zero-effect regions the sign of the marginal-median difference (MD-dMRI minus DTI) can differ from the paired Delta, and the paired Delta, $r_{rb}$, and q are the inferential quantities. q is the Benjamini-Hochberg false-discovery-rate-adjusted p-value within each (montage, Group 1) family; q < 0.001 is shown as <0.001.

| Montage | Region | ISO $p_{95}$ $|E|$ (V/m) | DTI (V/m) | MD-dMRI (V/m) | Delta (V/m) | 95% CI | $r_{rb}$ | q |
|---|---|---|---|---|---|---|---|---|
| M1 pad | Frontal cortex | 0.3731 | 0.3750 | 0.3730 | 0.0010 | [0.0006, 0.0020] | +0.65 | 0.003 |
| M1 pad | Parietal cortex | 0.2755 | 0.2733 | 0.2773 | 0.0032 | [0.0016, 0.0034] | +1.00 | <0.001 |
| M1 pad | Temporal cortex | 0.2397 | 0.2439 | 0.2435 | 0.0006 | [0.0000, 0.0009] | +0.43 | 0.048 |

| Montage | Region | ISO $p_{95}$ $\|E\|$ (V/m) | DTI (V/m) | MD-dMRI (V/m) | Delta (V/m) | 95% CI | $r_{rb}$ | q |
|---|---|---|---|---|---|---|---|---|
| M1 pad | Occipital cortex | 0.1690 | 0.1673 | 0.1676 | 0.0008 | [0.0000, 0.0011] | +0.51 | 0.019 |
| M1 pad | Frontal WM | 0.5114 | 0.5637 | 0.5341 | -0.0239 | [-0.0298, -0.0214] | -1.00 | <0.001 |
| M1 pad | Parietal WM | 0.3789 | 0.4137 | 0.4008 | -0.0163 | [-0.0197, -0.0146] | -1.00 | <0.001 |
| M1 pad | Temporal WM | 0.3278 | 0.3787 | 0.3580 | -0.0194 | [-0.0242, -0.0168] | -1.00 | <0.001 |
| M1 pad | Occipital WM | 0.2214 | 0.2417 | 0.2324 | -0.0110 | [-0.0127, -0.0091] | -1.00 | <0.001 |
| M1 pad | Corpus callosum | 0.4663 | 0.5010 | 0.4960 | -0.0053 | [-0.0077, -0.0008] | -0.59 | 0.007 |
| M1 pad | Mesencephalon | 0.3134 | 0.3185 | 0.3244 | 0.0005 | [-0.0021, 0.0044] | -0.02 | 0.949 |
| M1 pad | Pons | 0.2219 | 0.2397 | 0.2334 | -0.0050 | [-0.0078, -0.0018] | -0.86 | <0.001 |
| DLPFC pad | Frontal cortex | 0.3248 | 0.3221 | 0.3277 | 0.0047 | [0.0041, 0.0055] | +1.00 | <0.001 |
| DLPFC pad | Parietal cortex | 0.1109 | 0.1104 | 0.1109 | 0.0005 | [0.0001, 0.0009] | +0.54 | 0.014 |
| DLPFC pad | Temporal cortex | 0.1402 | 0.1421 | 0.1424 | 0.0008 | [0.0002, 0.0010] | +0.63 | 0.004 |
| DLPFC pad | Occipital cortex | 0.0729 | 0.0718 | 0.0723 | 0.0008 | [0.0005, 0.0009] | +0.88 | <0.001 |

| Montage | Region | ISO $p_{95}$ $\|E\|$ (V/m) | DTI (V/m) | MD-dMRI (V/m) | Delta (V/m) | 95% CI | $r_{rb}$ | q |
|---|---|---|---|---|---|---|---|---|
| DLPFC pad | Frontal WM | 0.4487 | 0.4927 | 0.4738 | -0.0220 | [-0.0279, -0.0195] | -1.00 | <0.001 |
| DLPFC pad | Parietal WM | 0.1573 | 0.1715 | 0.1657 | -0.0066 | [-0.0091, -0.0059] | -1.00 | <0.001 |
| DLPFC pad | Temporal WM | 0.1826 | 0.2065 | 0.1986 | -0.0082 | [-0.0106, -0.0072] | -1.00 | <0.001 |
| DLPFC pad | Occipital WM | 0.0992 | 0.1079 | 0.1036 | -0.0047 | [-0.0055, -0.0042] | -1.00 | <0.001 |
| DLPFC pad | Corpus callosum | 0.2794 | 0.3119 | 0.3097 | -0.0025 | [-0.0036, 0.0014] | -0.29 | 0.190 |
| DLPFC pad | Mesencephalon | 0.1714 | 0.1849 | 0.1879 | 0.0000 | [-0.0023, 0.0018] | -0.11 | 0.624 |
| DLPFC pad | Pons | 0.1447 | 0.1512 | 0.1509 | -0.0030 | [-0.0042, -0.0014] | -0.85 | <0.001 |
| HD-M1 | Frontal cortex | 0.2094 | 0.2077 | 0.2107 | 0.0028 | [0.0019, 0.0037] | +1.00 | <0.001 |
| HD-M1 | Parietal cortex | 0.2277 | 0.2285 | 0.2299 | 0.0013 | [0.0007, 0.0015] | +0.75 | <0.001 |
| HD-M1 | Temporal cortex | 0.1328 | 0.1324 | 0.1337 | 0.0023 | [0.0015, 0.0031] | +1.00 | <0.001 |
| HD-M1 | Occipital cortex | 0.0476 | 0.0474 | 0.0477 | 0.0006 | [0.0006, 0.0009] | +0.98 | <0.001 |
| HD-M1 | Frontal WM | 0.2705 | 0.2858 | 0.2805 | -0.0097 | [-0.0121, -0.0075] | -1.00 | <0.001 |

| Montage | Region | ISO $p_{95}$ $\|E\|$ (V/m) | DTI (V/m) | MD-dMRI (V/m) | Delta (V/m) | 95% CI | $r_{rb}$ | q |
|---|---|---|---|---|---|---|---|---|
| HD-M1 | Parietal WM | 0.2852 | 0.3069 | 0.2968 | -0.0105 | [-0.0134, -0.0081] | -1.00 | <0.001 |
| HD-M1 | Temporal WM | 0.1784 | 0.2055 | 0.1927 | -0.0114 | [-0.0128, -0.0093] | -1.00 | <0.001 |
| HD-M1 | Occipital WM | 0.0634 | 0.0660 | 0.0645 | -0.0013 | [-0.0021, -0.0011] | -1.00 | <0.001 |
| HD-M1 | Corpus callosum | 0.0894 | 0.0949 | 0.0952 | -0.0002 | [-0.0006, 0.0006] | -0.03 | 0.898 |
| HD-M1 | Mesencephalon | 0.0612 | 0.0656 | 0.0667 | -0.0002 | [-0.0008, 0.0012] | -0.03 | 0.898 |
| HD-M1 | Pons | 0.0459 | 0.0497 | 0.0478 | -0.0013 | [-0.0018, -0.0008] | -0.94 | <0.001 |
| HD-DLPFC | Frontal cortex | 0.2187 | 0.2166 | 0.2176 | 0.0012 | [0.0010, 0.0021] | +0.88 | <0.001 |
| HD-DLPFC | Parietal cortex | 0.0591 | 0.0590 | 0.0593 | 0.0002 | [-0.0001, 0.0005] | +0.47 | 0.034 |
| HD-DLPFC | Temporal cortex | 0.0603 | 0.0603 | 0.0610 | 0.0009 | [0.0006, 0.0011] | +0.94 | <0.001 |
| HD-DLPFC | Occipital cortex | 0.0193 | 0.0189 | 0.0193 | -0.0001 | [-0.0001, 0.0000] | -0.42 | 0.055 |
| HD-DLPFC | Frontal WM | 0.2828 | 0.3094 | 0.2966 | -0.0110 | [-0.0170, -0.0105] | -1.00 | <0.001 |
| HD-DLPFC | Parietal WM | 0.0877 | 0.0960 | 0.0909 | -0.0021 | [-0.0032, -0.0018] | -1.00 | <0.001 |

| Montage | Region | ISO $p_{95}$ $\|E\|$ (V/m) | DTI (V/m) | MD-dMRI (V/m) | Delta (V/m) | 95% CI | $r_{rb}$ | q |
|---|---|---|---|---|---|---|---|---|
| HD-DLPFC | Temporal WM | 0.0852 | 0.0938 | 0.0897 | -0.0046 | [-0.0053, -0.0034] | -1.00 | <0.001 |
| HD-DLPFC | Occipital WM | 0.0279 | 0.0302 | 0.0289 | -0.0015 | [-0.0017, -0.0012] | -1.00 | <0.001 |
| HD-DLPFC | Corpus callosum | 0.1719 | 0.1868 | 0.1847 | -0.0033 | [-0.0046, -0.0008] | -0.53 | 0.017 |
| HD-DLPFC | Mesencephalon | 0.0466 | 0.0491 | 0.0492 | 0.0000 | [-0.0004, 0.0009] | +0.12 | 0.559 |
| HD-DLPFC | Pons | 0.0414 | 0.0441 | 0.0429 | -0.0007 | [-0.0011, -0.0004] | -0.81 | <0.001 |

**S-Table 2.** Predicted electric-field exposure of the Group-2 deep nuclei under each of the four tDCS montages (M1 pad, DLPFC pad, HD-M1, HD-DLPFC), n = 29. Each cell is the cohort median, with the interquartile range across the 29 participants in brackets, of the within-subject 95th-percentile electric-field magnitude ($p_{95}$ $|E|$, V/m) under the MD-dMRI conductivity model, over the gray-matter and white-matter elements of the nucleus mask. The nine bilateral CIT168 nuclei (putamen, caudate, nucleus accumbens, external and internal globus pallidus, substantia nigra pars compacta and reticulata, red nucleus, and subthalamic nucleus; L and R hemispheres, 18 masks) are exploratory and field-only, with no cross-comparison against the cohort's mechanical or microstructural measures. They are resolution-limited: the basal ganglia (putamen, caudate, nucleus accumbens) are well-resolved (minimal overlap, modest partial volume), whereas the midbrain and subthalamic nuclei sit at or below the 2.5 mm diffusion voxel and are additionally subject to inter-nucleus overlap, so their values should be read as order-of-magnitude field exposure rather than precise per-nucleus estimates.

| Nucleus | M1 pad | DLPFC pad | HD-M1 | HD-DLPFC |
|---|---|---|---|---|
| Putamen (L) | 0.392 [0.343, 0.427] | 0.279 [0.265, 0.307] | 0.141 [0.124, 0.160] | 0.087 [0.079, 0.096] |
| Caudate (L) | 0.451 [0.380, 0.494] | 0.348 [0.304, 0.394] | 0.173 [0.149, 0.234] | 0.101 [0.084, 0.116] |
| Nucleus accumbens (L) | 0.285 [0.270, 0.316] | 0.234 [0.225, 0.249] | 0.060 [0.054, 0.064] | 0.080 [0.076, 0.089] |
| Globus pallidus ext (L) | 0.295 [0.258, 0.336] | 0.220 [0.198, 0.232] | 0.100 [0.084, 0.114] | 0.068 [0.064, 0.077] |

| Nucleus | M1 pad | DLPFC pad | HD-M1 | HD-DLPFC |
|---|---|---|---|---|
| Globus pallidus int (L) | 0.293 [0.256, 0.313] | 0.201 [0.179, 0.215] | 0.087 [0.080, 0.104] | 0.066 [0.060, 0.072] |
| Substantia nigra pc (L) | 0.290 [0.250, 0.327] | 0.153 [0.133, 0.168] | 0.046 [0.043, 0.059] | 0.036 [0.032, 0.039] |
| Substantia nigra pr (L) | 0.283 [0.252, 0.337] | 0.157 [0.142, 0.188] | 0.060 [0.054, 0.063] | 0.041 [0.038, 0.046] |
| Red nucleus (L) | 0.352 [0.319, 0.412] | 0.176 [0.157, 0.209] | 0.065 [0.056, 0.070] | 0.042 [0.037, 0.047] |
| Subthalamic nucleus (L) | 0.249 [0.228, 0.303] | 0.145 [0.133, 0.173] | 0.060 [0.056, 0.067] | 0.047 [0.039, 0.054] |
| Putamen (R) | 0.350 [0.318, 0.377] | 0.313 [0.276, 0.336] | 0.051 [0.047, 0.058] | 0.070 [0.063, 0.078] |
| Caudate (R) | 0.389 [0.348, 0.434] | 0.345 [0.286, 0.378] | 0.062 [0.054, 0.072] | 0.074 [0.062, 0.079] |
| Nucleus accumbens (R) | 0.298 [0.277, 0.345] | 0.246 [0.231, 0.281] | 0.046 [0.041, 0.050] | 0.068 [0.060, 0.073] |
| Globus pallidus ext (R) | 0.327 [0.304, 0.339] | 0.273 [0.253, 0.294] | 0.055 [0.050, 0.060] | 0.067 [0.062, 0.074] |
| Globus pallidus int (R) | 0.268 [0.253, 0.300] | 0.199 [0.190, 0.228] | 0.047 [0.041, 0.051] | 0.054 [0.050, 0.059] |
| Substantia nigra pc (R) | 0.228 [0.207, 0.245] | 0.138 [0.123, 0.154] | 0.036 [0.034, 0.040] | 0.034 [0.031, 0.036] |
| Substantia nigra pr (R) | 0.244 [0.233, 0.267] | 0.156 [0.141, 0.175] | 0.038 [0.037, 0.043] | 0.036 [0.032, 0.040] |
| Red nucleus (R) | 0.322 [0.266, 0.358] | 0.172 [0.144, 0.193] | 0.060 [0.045, 0.071] | 0.040 [0.033, 0.051] |
| Subthalamic nucleus (R) | 0.264 [0.241, 0.280] | 0.171 [0.156, 0.186] | 0.048 [0.044, 0.052] | 0.043 [0.039, 0.052] |

**S-Table 3.** Per-region orientation divergence between the QTI mean tensor  and the single-shell DTI tensor, n = 29. V1 angle (own registration) is the cohort median of each subject's median acute angle between the two principal eigenvectors over positive-definite voxels in the region, with each tensor carried to T1 by its own 12-DOF affine; V1 angle (shared registration) is the same quantity after both tensors are carried to T1 through a single shared registration (the matched-registration control shown in Figure 5). ΔFA is the cohort-median fractional-anisotropy difference ( minus DTI; positive =  more anisotropic) and $\Delta(\lambda_1/\lambda_3)$ the corresponding largest-to-smallest eigenvalue-ratio difference; both are preprocessing-sensitive magnitude descriptors, reported for completeness rather than as findings. Ctx, cortical lobe; WM, white-matter lobe; CC, corpus callosum. The principal-direction angle is shown in Figure 5.

| Region | V1 angle, own reg. (deg) | V1 angle, shared reg. (deg) | ΔFA | $\Delta(\lambda_1/\lambda_3)$ | n |
|---|---|---|---|---|---|

| Region | V1 angle, own reg. (deg) | V1 angle, shared reg. (deg) | ΔFA | $\Delta(\lambda_1/\lambda_3)$ | n |
|---|---|---|---|---|---|
| WM Frontal | 18.2 | 20.1 | -0.06 | -0.21 | 29 |
| WM Parietal | 18.1 | 18.9 | -0.08 | -0.27 | 29 |
| WM Temporal | 23.5 | 24.8 | -0.05 | -0.15 | 29 |
| WM Occipital | 25.8 | 26.2 | -0.06 | -0.16 | 29 |
| Corpus callosum | 12.8 | 12.8 | -0.04 | -0.07 | 29 |
| Mesencephalon | 18.0 | 17.7 | +0.06 | +0.25 | 29 |
| Pons | 14.8 | 16.1 | -0.01 | +0.00 | 29 |
| Ctx Frontal | 43.8 | 44.4 | +0.00 | +0.00 | 29 |
| Ctx Parietal | 43.5 | 44.2 | +0.00 | +0.00 | 29 |
| Ctx Temporal | 45.8 | 45.9 | +0.02 | +0.06 | 29 |
| Ctx Occipital | 47.2 | 46.8 | +0.03 | +0.07 | 29 |
